# Dopamine and cortical iPSC-derived neurons with different Parkinsonian mutations show variation in lysosomal and mitochondrial dysfunction: implications for protein deposition versus selective cell loss

**DOI:** 10.1101/2024.10.07.617117

**Authors:** Jessica Chedid, Yan Li, Adahir Labrador-Garrido, Dad Abu-Bonsrah, Chiara Pavan, Tyra Fraser, Dmitry Ovchinnikov, Melanie Zhong, Ryan Davis, Dario Strbenac, Jennifer A. Johnston, Lachlan H. Thompson, Deniz Kirik, Clare L. Parish, Glenda M. Halliday, Carolyn M. Sue, Gautam Wali, Nicolas Dzamko

## Abstract

**Background:** Mutations causing Parkinson’s disease (PD) give diverse pathological phenotypes whose cellular correlates remain to be determined. For example, those with *PRKN* loss of function mutations have significantly earlier selective vulnerability of dopamine neurons, those with *SNCA* mutations have increased alpha-synuclein deposition, while those with *LRRK2* mutations have additional deposition of tau. Yet all three mutation types are implicated in mitochondrial and/or lysosomal dysfunction. Direct comparison of cell models with these mutations would clarify the relative cellular dysfunctions associated with these different pathological phenotypes.

**Methods:** An unbiased high-content imaging platform using orthogonal probes to assess both lysosomal and mitochondrial dysfunction, along with alpha-synuclein and tau protein deposition was established using induced pluripotent stem cell (iPSC) derived cortical and ventral midbrain neurons. Three mutation types, *SNCA* A53T, *LRRK2* R1441G and *PRKN* loss of function (lof), were selected as exemplars of divergent PD pathological phenotypes and compared to each other, and to control iPSC from subjects without PD.

**Results:** Different PD mutations caused cell type specific dysfunctions, likely to impact on both selective neuronal vulnerability and the pathologies observed in PD. Comparison of dopamine neurons identified that both lysosomal and mitochondrial dysfunction were predominant with *PRKN* lof mutations, whereas immunofluorescent staining revealed that *SNCA* A53T and *LRRK2* R1441G mutations had increased tau deposition. In contrast, cortical neurons with *SNCA* and *LRRK2* mutations both had mitochondrial and autophagy impairments without protein deposition, with *LRRK2* cells additionally showing decreased glucocerebrosidase activity and increased alpha-synuclein phosphorylation.

**Conclusions:** Lysosomal and mitochondrial dysfunction are predominant in dopamine neurons with *PRKN* lof mutations, and may drive the early selective loss of dopamine neurons in *PRKN* mutation carriers. More subtle cellular abnormalities in the *SNCA* A53T cell lines are likely to predispose to alpha-synuclein aggregation and tau protein deposition over time. The *LRRK2* R1441G may also predispose to tau deposition, but despite substantial lysosomal dysfunction with increased alpha-synuclein phosphorylation, pathological alpha-synuclein accumulations were not observed. Understanding the mechanistic differences in how lysosomal and mitochondrial dysfunction impact on PD pathogenesis in different disease subtypes may be important for therapeutic development.

## Background

Parkinson’s disease (PD) is the most common neurodegenerative movement disorder, with its classic motor symptoms the result of progressive loss of dopamine producing neurons in the midbrain, while broader neuronal dysfunction and symptomology are associated with pathological protein depositions of misfolded alpha-synuclein into cytoplasmic inclusions termed Lewy bodies [1]. The spreading of pathological alpha-synuclein through the brain in PD largely follows a defined staging pattern from brainstem through to neocortical regions, associating with an array of clinical symptoms including autonomic dysfunction, REM sleep disorder, depression and cognitive decline [2, 3]. The presence of co-pathologies in PD brain, for example hyperphosphorylation and/or abnormal deposition of the protein tau, can further modify the disease course and contribute to heterogenous symptomology [4]. As recent antibody trials targeting alpha-synuclein have not proved efficacious [5, 6], there is an unmet need to further understand the diversity of cellular mechanisms involved in pathogenesis of PD.

Diverse mechanisms associated with PD have become apparent due to the substantial genetic heterogeneity underlying a greater risk for or causing PD (associated with environmental circumstances) [7]. This molecular diversity associates with certain PD phenotypes. For example, *SNCA* gene mutations and multiplications invariably have a more aggressive disease that associates with widespread pathological alpha-synuclein deposition [8], while *PRKN* loss of function gene abnormalities have early-onset, focal degeneration of long duration with no or limited spread of pathological alpha-synuclein [9]. Mutations in *LRRK2* on the other hand, give a typical late-onset disease, but with variable combinations of underlying alpha-synuclein and/or tau pathology [10-13]. These distinct genetic phenotypes can be modelled in the laboratory to determine what cellular mechanisms underlie the pathological diversity observed in PD.

Many studies have used neurons differentiated from patient derived induced pluripotent stem cells (iPSC) to determine cellular dysfunction in PD. Main overlapping cellular mechanisms of dysfunction across *SNCA*, *PRKN* and *LRRK2* genotypes occur within the mitochondrial and autophagy-lysosomal pathways [14-16]. However, it is still unclear if some mutations drive cellular phenotypes to a greater extent than others, as side-by-side comparisons across different mutation groups are rarely performed. Comparative genotype studies using differentially susceptible cell types are even fewer. There are technical reasons for this, particularly the challenge of ensuring scale and diversity in the populations of cells being compared to model typical human populations. For example, a review of 67 recent iPSC PD studies indicated that the average number of cell lines used in a study was five (comprised of comparing two control lines to three disease lines, usually harbouring the same gene mutation) [17]. At least one study has investigated selective vulnerability by comparing dopamine and cortical neurons differentiated from the same iPSC lines containing duplication of *SNCA*, demonstrating increased alpha-synuclein aggregation and oxidative stress selectively in dopamine neurons [18]. Moreover, mitochondrial morphology deficits were present in iPSC-derived dopaminergic neurons with the *LRRK2* R1441C mutation, but not observed in cortical neurons with the same mutation from rodent cultures [19]. These studies clearly indicate that neuron-type specific effects of PD-associated mutations are important to define, but to compare such phenotypes across different mutation types require advances in high throughout imaging protocols and cell line availability.

To further advance knowledge on the extent of cellular dysfunction across different mutation groups in different PD relevant neuron types, we have established an unbiased high-content imaging platform with orthogonal probes to assess lysosomal dysfunction along with alpha-synuclein and tau protein deposition in iPSC derived cortical and ventral midbrain dopamine neurons compared to controls (n=3-4) without PD or PD gene mutations (n=3 cell lines per genotype). Mitochondrial respiration was also assessed across the mutation groups and cell types. We have focussed on two aspects of the disease – 1) dysfunction that may underlie the early selective vulnerability of dopamine cell loss in those with *PRKN* loss of function mutations compared with the later loss in the other genotypes, and 2) dysfunction that may underlie the abnormal protein deposition which is substantial in both dopamine and cortical neurons with *SNCA* triplications, and also observed in *LRRK2* R1441 mutations but rarely in *PRKN* loss of function mutations (particularly in cortical neurons). Our results indicate that *PRKN* mutations impact upon mitochondrial function in both neuron types, but lysosomal function and alpha-synuclein pathology are particularly impacted in dopamine neurons vulnerable to cell loss. *LRRK2* and *SNCA* mutations resulted in tau pathology in dopamine neurons, however mitochondrial and lysosomal dysfunction were more prevalent in cortical cells with the *LRRK2* R1441G mutation. Thus, the extent of cellular dysfunction associated with PD mutations may differ between neuron types, which has implications for understanding different aspects of PD pathophysiology related to dopamine neuron loss and the propagation of pathological proteins, as well as how PD may be treated.

## Methods

### Cell lines

Induced pluripotent stem cell (iPSC) lines with *LRRK2* R1441G (n=3) and *SNCA* A53T (n=3) used in the analyses presented in this article were obtained from the Golub Capital iPSC Parkinson’s Progression Markers Initiative (PPMI) Sub-study (https://www.ppmi-info.org/access-data-specimens/request-cell-lines). As such, the investigators within PPMI contributed to the design and implementation of PPMI and/or provided data and collected samples but did not participate in the analysis or writing of this report. For up-to-date information on the study, visit PPMI-info.org. mutations were obtained from the Golub Capital iPSC Parkinson’s Progression Marker Initiative (PPMI) sub-study (www.ppmi-info.org/cell-lines). Neurologically normal control lines (n=2) were also obtained from PPMI. The control line KOLF2-1J was obtained from the Jackson Laboratory (#JIPSC1000) and has been described previously [20]. The in-house control line RM3.5 has also been described previously [21, 22]. *PRKN* mutation iPSCs (n=3) were reprogrammed from fibroblasts in house, with the characterisation of one iPSC line reported [23]. Publications describing the characterisation of the remaining two *PRKN* mutation iPSC lines are pending. Available demographic data accompanying the iPSCs are shown in Table S1. Quality control experiments performed on the newly generated PRKN iPSC lines are shown in Table S2. The genotype of PPMI lines was confirmed by sequencing. Experiments on human iPSCs were approved by the Human Research Ethics Committee at the University of Sydney (2017/094).

### Ventral midbrain dopamine differentiation

Ventral midbrain (VM) progenitors were generated according to the methods of Gantner and colleagues [24]. At day 19 of differentiation, VM progenitors were cryopreserved in freeze media containing 20% DMSO, 60% KnockOut serum replacement and 20% media comprised of a 1:1 mix of Dulbecco’s Modified Eagle Medium (DMEM) F12 and neurobasal media with 2% B27 + vitamin A, 1% each N2 supplement, non-essential amino acids, ITS-A and glutamax and 0.5% penicillin/streptomycin (all from Thermofisher) and supplemented with 20 ng/mL BDNF (R&D), 20 ng/mL GDNF (R&D), 0.1 mM dibutyryl cAMP (Stem Cell Technologies), 200 nM ascorbic acid (Stem Cell Technologies), 1 ng/mL TGFβ3 (Lonza), 10 mM DAPT (Sigma) and 10 mM ROCK inhibitor Y27632 (Stem Cell Technologies) (subsequently referred to as ‘maturation media’). Cryovials were retained in vapour phase liquid nitrogen until required. Progenitors were thawed in maturation media and plated onto PhenoPlate 96 well plates (Revvity) pre-coated with 10 mg/mL human Laminin-521 (BioLamina), at a density of 30,000 cells per well. Cells were cultured in maturation media for a further 21 days, with 90% of the media replaced every second day with fresh supplemented maturation media. At day 21 post-thaw, a total of 40 days of differentiation, the cells were used for experiments. Differentiated VM neurons from all lines were stained with forkhead box protein A2 (FOXA2) and tyrosine hydroxylase (TH) to confirm the presence of dopamine neurons.

### Cortical neuron differentiation

Cortical neural progenitors were generated from iPSC according to the methods of Gantner and colleagues [25]. At day 25 of differentiation, vials of cortical progenitors were cryopreserved and retained in a vapour phase liquid nitrogen tank until required. Progenitors were thawed onto PhenoPlate 96 well plates (Perkin Elmer) pre-coated with 15 µg/ml poly-L-ornithine (Sigma) and 10 µg/ml mouse laminin (Sigma), at a density of 10,000 cells per well in the presence of 10 µM Y27632. Cells were initially thawed and cultured in cortical base media comprised of a 1:1 mix of DMEM F12 and neurobasal media containing 0.5 x B27 supplement, 0.5 x N2 supplement, 0.5 x ITS-A, 0.5 x non-essential amino acids, 0.5 x glutamax, 50 U/ml penicil/streptomycin and 49.5 μM β-mercaptoethanol (all from Thermofisher). Media was changed on day 1 to remove the ROCK inhibitor Y27632. From day 3, the media was changed to a cortical maturation media comprised of a 1:1 mix of DMEM F12 and neurobasal media containing 1x B27 supplement, 1x N2 supplement, 1x ITS-A, 1x non-essential amino acids, 0.5x glutamax and 50 U/ml penicil/streptomycin and further supplemented with 40 ng/ml BDNF, 40 ng/ml GDNF (both Stem Cell Technologies), 50 μM dibutyryl cAMP (Stem Cell Technologies), 200 nM ascorbic acid (Sigma), 100 ng/ml mouse laminin and 10 μM of the γ-secretase inhibitor DAPT (Sigma) as previously optimised [26]. Cells were differentiated for 12 days in cortical maturation media, with 90% of the media replaced every second day with fresh media. At day 15 post-thaw, a total of 40 days of differentiation, the cells were used for experiments. Differentiated cortical cells from all lines were stained with POU domain transcription factor 2 (BRN2) to identify cortical layers II/III, COUP-TF-interacting protein 2 (CTIP2) to identify layer V cells, T-box brain protein 1 (TBR1) to identify deep-layer VI cells and microtubule-associated protein 2 (MAP2) as a general neuronal marker.

### Fixed cell staining

For fixed staining experiments, half of the media was discarded from the wells and replaced with the same volume of 4% paraformaldehyde (PFA) and incubated at room temperature in the dark for 10 min. Then the mixture of cell culture media and PFA was gently removed and replaced with 4% PFA and incubated for a further 15 min. PFA was then removed, and the wells gently washed with 1 x phosphate buffered saline (PBS). Cells were then permeabilized with 0.3% Triton X-100 in 1 x PBS for 20 min, blocked in 3% bovine serum albumin (BSA) (Bovogen) with 0.1% Triton-X in 1 x PBS for 1 h, and incubated overnight at 4 °C in primary antibody in blocking solution. After overnight incubation, cells were washed 3 × 5 min in 1 × PBS and incubated with secondary antibodies diluted in blocking solution for 1 h at room temperature in the dark. Cells were washed again for 3 × 5 min in 1 × PBS with DAPI added to the last wash at a concentration of 1:10,000. Plates were stored in the dark at 4 °C until imaged. Primary and secondary antibodies used for fixed cell imaging are provided in Table S3 and Table S4 respectively. For the study of autophagic flux, half of the wells were pre-treated with 400 nM bafilomycin A1 (Sigma) for 4 h prior to fixing.

### Live cell staining

Multiple live cell assays were performed for lysosomal phenotyping. Plates were imaged in the Opera Phenix at 37°C and 5% CO2 (in air), which were set 30 min prior to the start of imaging. Working dilutions of the live cell imaging reagents were prepared in media and added to the wells (after discarding old media) before imaging. Live cell stains employed in this study are detailed in Table S5. DQ red BSA was imaged 90 min after the addition of the probe. For the glucocerebrosidase (GCase) substrate, 5-(pentafluorobenzoylamino)fluorescein di-ß-D-glucopyranoside (PFB-FDGlu), half of the wells were pre-treated with either 0.5 mM GCase inhibitor conduritol-β-epoxide (CBE) dissolved in DMSO, or DMSO alone (vehicle) for 1 h before adding PFB-FDGlu at 0.375 mM. Cells were imaged at 15, 30 and 45 min to measure GCase substrate fluorescence over time. The GCase activity index was calculated as the ratio of PFB-FDGlu MFI -CBE / PFB-FDGlu + CBE as has been previously established [27].

### High content imaging

Images were acquired on the Opera Phenix high-content system with either 20x or 40x water objective and analysed using Harmony software (Revvity). Channels were chosen for imaging depending on the combination of antibodies or probes used for live or fixed imaging and best available channels to avoid spectral overlap. Z-stacks were acquired with a set number of Z-planes and Z-steps. Details on the magnification, average number of cells imaged, channels used, number of fields of view acquired, Z-steps and number of planes in each Z-stack for each acquired variable are shown in Table S6 (cortical neurons) and Table S7 (VMDA neurons). For the experiments using live cell imaging the number of Z-steps acquired per stack was kept low to ensure the entire experiment could be images in less than 30 min with consistent results.

### High content image analysis

Images were analysed using Harmony software (Revvity) with flatfield and brightfield corrections applied and stack processing set to maximum projection. DAPI or Hoechst were used to identify nuclei, with nuclei morphology settings applied to eliminate non-specific signals, dead cells, and large clumps of unsegmented nuclei. Nuclei touching the borders of the image were also excluded. Cytopainter green or MAP2 or TH were used to create masks for cells of interest. The median fluorescent intensity (MFI) was determined for regions of interest using the Intensity Properties building block. Puncta were detected using the Find Spots building block, and the size and area of puncta determined with the Morphology building block applied to the spots. To optimize spot detection, detection sensitivity, splitting coefficient and background correction parameters were manually tuned and applied across different cell lines within each staining experiment. Detected spots could then be filtered by intensity and morphology properties. Visual inspection of the images was used to verify and fine tune analysis sequences across independent differentiation rounds for higher consistency. Data were exported to Excel and then to SPSS for analysis.

### Immunoblotting

Cortical cells were lysed in buffer comprised of CelLytic TM cell lysis buffer (Sigma) with 100X Halt Protease Inhibitor Cocktail (ThermoFisher). Lysates were centrifuged at 13,000 x g at 4℃ for 20 min and protein concentrations in supernatant were determined by bicinchoninic acid assay (Thermofisher) as per the manufacturer instructions. Cell lysate was then mixed with sample buffer (2% SDS, 20% glycerol, 2.5% bromophenol blue, 12.5 mM Tris-HCl, pH 6.8, 5% 2-mercaptoethanol) and heated at 70℃ for 10 min before separating on SDS-PAGE gels. 20 µg of protein was loaded per well on Nu-Page 4-12% Bis-Tris protein gels (ThermoFisher) for each experiment. The electrophoresis took approximately 45-60 min at 150-180 volts. Precision Plus Protein Dual Colour Standards (Bio-Rad) were used to determine molecular weights of target bands. After separation, proteins were transferred to 0.45 μm nitrocellulose membrane (Bio-Rad) at 90 volts for 90 min by a wet transfer system (Bio-Rad) in the presence of transfer buffer (0.14 M glycine, 186 mM Tris, 15% methanol). Membranes were blocked in 5% (w/v) skim milk or 5% (w/v) BSA dissolved in 1 x TBST (0.87% NaCl, 0.01 M Tris, pH 7.4, 0.05% Tween-20) solution for 1 h, followed by incubation with primary antibodies at 4℃ overnight and in horseradish peroxidase (HRP)-conjugated secondary antibodies at room temperature for 2 h. Primary and secondary antibodies were diluted in 5% and 2.5% (w/v) skim milk or BSA/TBST solution, respectively. Three washing steps with 1x TBST buffer were applied after the incubation of each antibody, 10 min per wash. Primary antibodies used for immunoblot in this study are shown in Table S3. Bands were visualized in a Bio-Rad Chemidoc MP system after evenly covering the blot with SuperSignal West Femto Maximum Sensitivity Substrate (Thermofisher). The intensities of protein bands were quantified using ImageLab software (Bio-Rad). Results are shown as arbitrary units normalized to β-actin.

### Extracellular oxygen consumption assay

Extracellular oxygen consumption was measured using an oxygen-sensitive fluorescent dye (Extracellular Oxygen Consumption Assay Kit, Abcam) as per the manufacturer protocol. 1 x 10^6^ progenitor cells were seeded in each well of a 96 well plate in culture media and differentiated as above. On the day of analysis, the culture media was replaced with the assay media (oxygen-sensitive fluorescent dye diluted in culture media at 1:15 dilution). A layer of mineral oil was then added on top of the assay media using a dropper to restrict the diffusion of oxygen. The fluorescence signal was measured using a Nivo microplate reader (Revvity) at 37°C for 120 min with readings at 2-minute intervals at wavelengths of 340 nm for excitation and 650 nm for emission. Fluorescent intensity values were normalised to the 30 min time, serving as a reference point. The temporal changes in fluorescence intensity were depicted using XY graphs in GraphPad Prism. To measure the rate of change, a linear segment of the graph was identified for each graph and a simple linear regression model was fitted to this segment to calculate the slope for each PD mutation, representing the rate of change in fluorescence intensity over time. The slopes or regression coefficients of PD groups were compared with the control using one-way ANOVA, and their statistical significance levels are marked in graphs. Experiments were repeated at least three times.

### Data analysis

Data analysis was performed using SPSS (IBM) or Prism (Graphpad) with graphs made using Prism. All experiments were performed with at least 2 independent rounds of differentiation and at least in duplicate. In most cases, 3 independent rounds of differentiation each in triplicate was performed. For characterisation, lysosomal and pathological measures, analysis of variance, covarying for differentiation round and employing Bonferroni corrected post-hoc testing was used to determine group effects for each variable. Significance was accepted at P < 0.05. Kolmogorov-Smirnov testing was used to determine normal distribution, with non-normal data then either log or square root transformed to achieve normality. SPSS was used to remove statistical outliers. Missing data were mean imputed. Less than 1% of the data set comprised outliers or missing data. Mitochondrial oxygen consumption was analysed with Prism software using One-Way ANOVA and Tukey post hoc test.

## Results

### Successful differentiation of ventral midbrain dopamine and cortical neurons with SNCA, LRRK2 and PRKN mutations

To determine if the impact of PD mutations on lysosomal and mitochondrial function varies with mutation or neuron type, cell lines were differentiated to ventral midbrain dopamine, and cortical neurons. At Day 40 of ventral midbrain neuronal differentiation, >70% of cells expressed the ventral floorplate transcription factor FOXA2, reflective of appropriate regional specification (Fig. 1A), with no significant differences in % of FoxA2+ cells between groups (Fig. 1B). Dopamine neurons were identified as TH-immunoreactive cells within the cultures (Fig. 1C), with a significantly higher number of dopamine neurons present in cultures derived from *SNCA* A53T mutation lines compared to all lines, and a higher number in the *LRRK2* R1441G mutation group compared to controls (Fig. 1D). The TH cell area was also increased in these two mutation groups (Supplementary Fig. 1A). For all differentiation measures, *PRKN* lof cells were indistinguishable from controls, however the overall cell count based on Hoechst nuclei staining was consistently lower in the *PRKN* lines (Supplementary Fig. 1B). These results show the successful differentiation to dopamine neurons for all cell lines, with higher numbers of TH cells present in *SNCA* A53T and *LRRK2* R1441G mutation cell lines.

**Figure 1.**
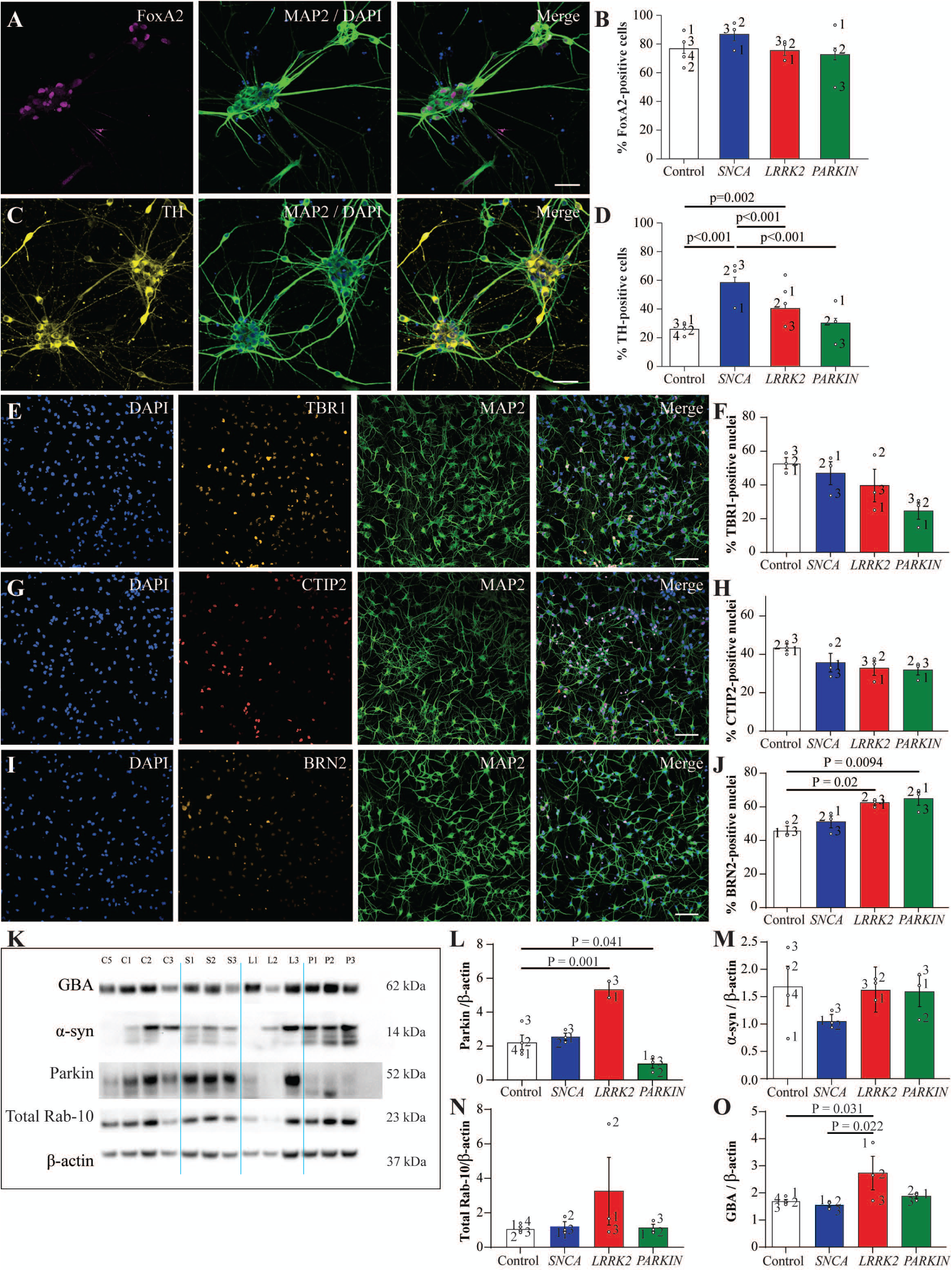
Successful differentiation of ventral midbrain dopamine and cortical neurons with *SNCA, LRRK2* and *PRKN* mutations. **A**) Representative images of VMDA neurons immunostained with FoxA2 (magenta), MAP2 (green) and DAPI (blue). **B**) Percentage of VMDA neurons expressing FoxA2 in the different mutation groups. **C**) Representative images of VMDA neurons immunostained with TH (yellow), MAP2 (green) and DAPI. **D**) Percentage of VMDA neurons expressing TH in the different mutation groups. Representative images of cortical neurons immunostained with TBR1 (orange), MAP2 (green) and DAPI (blue) (**E**) and subsequent quantification (**F**). Representative images of cortical neurons immunostained with CTIP2, MAP2 (green) and DAPI (blue) (**G**) and subsequent quantification (**H**). Representative images of cortical neurons immunostained with BRN2, MAP2 (green) and DAPI (blue) (**I**) and subsequent quantification (**J**). The scale bar in all images is 200 µm. Image quantification graphs show mean ± SEM. Each datapoint represents the mean value for each cell line derived from n=3 biological replications (each performed in at least duplicate). Representative Immunoblots (**K**) and subsequent quantification of Parkin (**L**), α-syn (**M),** Rab-10 (**N**) and glucocerebrosidase (**O**) proteins respectively in cortical neurons of different mutation groups. Protein expression is normalised to β-actin. Graphs show mean ± SEM with data points derived from n=2 independent experiments.

By day 40 of cortical differentiation, cells expressed markers indicative of deep layer VI (TBR1, Fig. 1E) and Layer V cortical neurons (CTIP2, Fig. 1G) as well as upper layer II-IV neurons (BRN2, Fig. 1I). Despite some subtle variation between individual lines, there was no significant effect of the mutations on the expression of the deep layer cortical differentiation markers (Fig. 1F and H), yet BRN2+ cells were significantly higher in *LRRK2* R1441G and *SNCA* A53T mutation lines compared to the controls (Fig. 1J). As more cortical cells were available, protein lysates could also be generated for additional western blot characterization to screen for proteins implicated in PD pathology or broader cell death cascades (Fig. 1K) (Fig. 1K). *PRKN* lof cell lines had significantly reduced levels of parkin protein compared to controls, whereas *LRRK2* R1441G mutation cell lines showed higher levels of Parkin (Fig. 1L). In contrast, alpha-synuclein levels were similar across groups, including those with *SNCA* A53T mutations (Fig. 1M). Total levels of Rab10 were similar across groups (Fig. 1N), however robust signal for RAB10 phosphorylated at the LRRK2 Thr73 site was not detected, noting that phosphatase inhibitors where not included in the cell lysis buffer. *LRRK2* R1441G cell lines did have an increase in the levels of glucocerebrosidase protein compared to other genotypes (Fig. 1O).

### PRKN and LRRK2 mutations are associated with impaired glucocerebrosidase (GCase) activity in dopamine and cortical neurons respectively

Lysosomal function was measured in the ventral midbrain and cortical cultures using live and fixed imaging. In the ventral midbrain differentiated neurons, lysosomal GCase activity index was significantly reduced in *PRKN* lof cells (Fig. 2A and B), whereas the accumulation of DQ red BSA (used to study endocytic trafficking) was unchanged between genotypes (Fig. 2C and D). In the cortical cells there was a significant effect of genotype, with lysosomal GCase activity index being lower in the *LRRK2* R1441G mutation group (Fig. 2E and F). As levels of GCase protein were available for cortical cell lines, the GCase index was also normalized to levels of GCase protein, which showed an even greater effect of the *LRRK2* mutation to reduce GCase activity (Fig. 2G). In the cortical cells, *SNCA* A53T mutations had a greater intensity of DQ red BSA signal compared to *LRRK2* R1441 mutations (Fig. 2H and I). This data shows that PD mutations affect lysosomal activity in a cell and mutation type specific manner, with *PRKN* lof mutations reducing GCase activity only in ventral midbrain neurons, whereas the *LRRK2* R1441G mutation reduced GCase activity only in cortical neurons.

**Figure 2.**
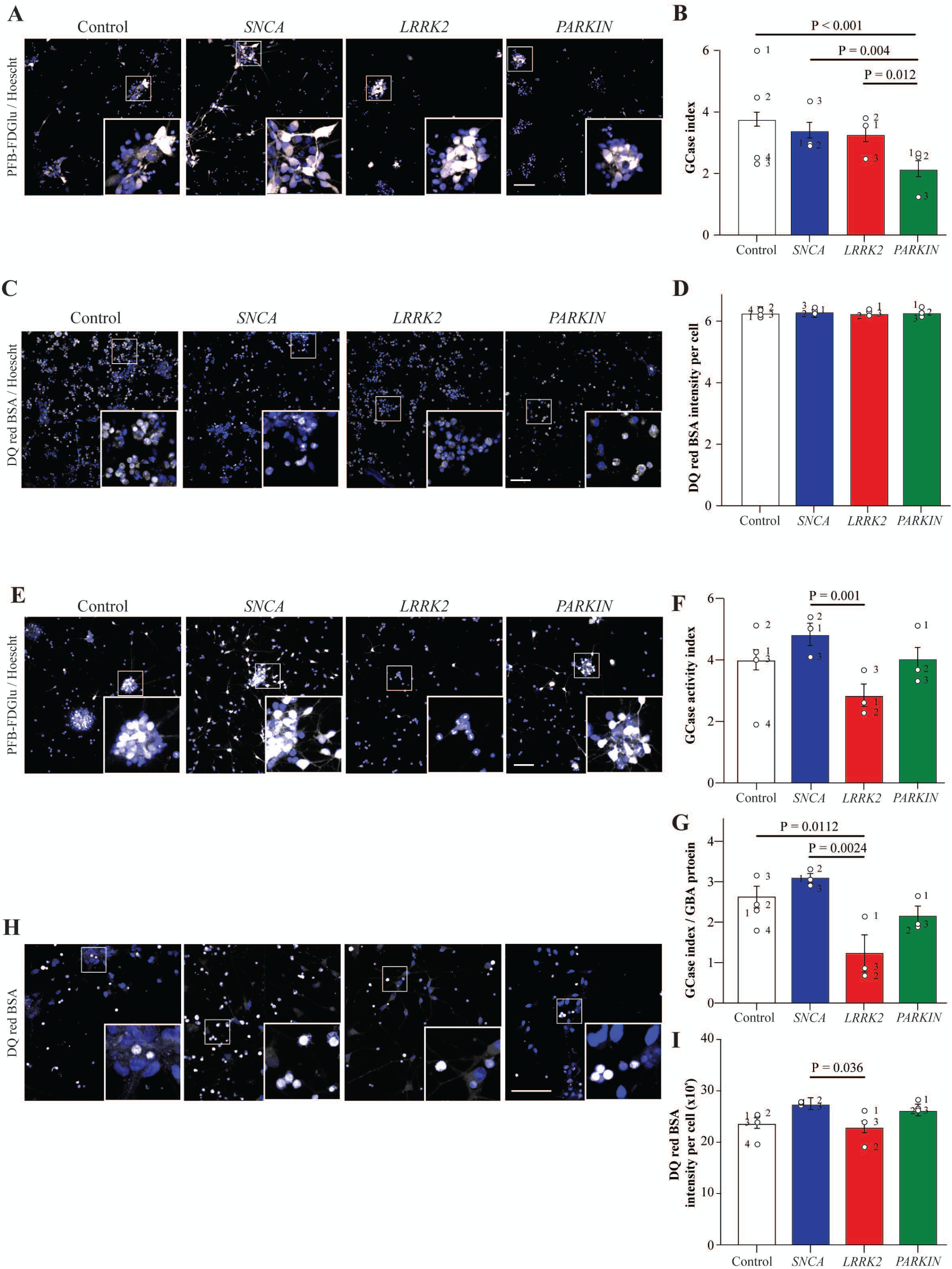
Glucocerebrosidase activity in VMDA and cortical neurons of the different mutation groups. **A**) Representative images of the PFB-FDGlu fluorescence (white) in VMDA neurons at 45 min after addition of the substrate, with Hoechst (blue). **B**) GCase activity index calculated for each cell line of the different mutation groups in VMDA neurons. **C**) Representative images of the DQ red BSA fluorescence (magenta) in VMDA neurons at 90 min after addition of the substrate with Hoechst (Blue). **D**) Intensity of DQ red BSA fluorescence per cell calculated for each cell line of the different mutation groups in VMDA neurons. **E**) Representative images of the PFB-FDGlu fluorescence (white) in the cortical neurons at 45 min after addition of the substrate, with Hoechst (blue). **F**) GCase activity index calculated for each cell line of the different mutation groups in cortical neurons. **G**) GCase activity index normalised with the protein expression level of GBA for each cell line of the different mutation groups. **H**) Representative images of the DQ red BSA fluorescence (white) in cortical neurons at 90 min after addition of the substrate with Hoechst (Blue). **I**) Intensity of DQ red BSA fluorescence per cell calculated for each cell line of the different mutation groups in cortical neurons. The scale bar in all images is 200 µm. Image quantification graphs show mean ± SEM. Each datapoint represents the mean value for each cell line derived from n=3 biological replications (each performed in at least duplicate).

### PRKN and SNCA mutations are associated with altered autophagy in dopamine and cortical neurons respectively

To assess lysosome-autophagy function in the dopamine and cortical neurons using fixed cell imaging, the lysosomal dysfunction marker galectin 3 (Gal3), the chaperone mediated autophagy marker LAMP2A, and the lysosomal biogenesis marker TFEB were assessed. As a measure of autophagy flux, the number of puncta per cell of the autophagy marker p62 was also measured, in the presence and absence of the autophagy inhibitor bafilomycin. For the TH+ midbrain differentiated cells, in the absence of bafilomycin, the number of p62 puncta per cell was significantly elevated in *LRRK2* R1441G mutation cells (Fig. 3A and B). Whilst in the presence of bafilomycin, the number of p62 puncta per cell was significantly reduced in the *PRKN* lof lines (Fig. 3A and C). When assessed as a ratio of +/- bafilomycin treatment, indicative of autophagic flux, the *LRRK2* R1441G mutation cells showed an impaired upregulation of p62 in response to bafilomycin compared to the control cells (Supplementary Fig 2A). There was no significant difference between the groups for the number per cell or size of Gal3 puncta in the dopamine neurons (Supplementary Fig. 2C-E). The number of LAMP2A puncta were also similar across groups in the midbrain dopamine neurons (Supplementary Fig. 2F and G), although *SNCA* A53T and *PRKN* lof cells had smaller LAMP2A puncta compared to control cells (Supplementary Fig. 2F and H). For the cortical differentiated cells, the number of p62 puncta per cell was significantly increased in the *SNCA* A53T mutation cells suggesting impaired baseline autophagy. In contrast, levels of p62 were significantly decreased compared to controls in the *LRRK2* R1441G mutation group (Fig. 3D and E). The same results for p62 were observed following treatment with bafilomycin (Fig. 3D and F). Confirming these findings, immunoblotting for p62 in the absence of bafilomycin also showed a significant increase in the *SNCA* A53T mutation cells (Fig. 3G and H). When assessed as a ratio of +/- bafilomycin treatment, *PRKN* mutation lines showed a greater autophagic flux than other cell lines (Supplementary Fig 2B). The number of Gal3 puncta per cell was significantly increased in the cortical neuron *SNCA* A53T group (Fig. 3I and K), while the size of the Gal3 puncta was increased in the *LRRK2* R1441G mutation group (Fig. 3I and L). The intensity of TFEB staining in the nucleus was also increased in the *SNCA* A53T mutation group (Fig. 3J and M). This data shows that PD mutations affect autophagy in a cell and mutation specific manner, with *PRKN* lof and *LRRK2* R1441G impairing autophagy only in dopamine neurons, whereas *SNCA* A53T mutations had impaired autophagy only in cortical neurons. In cortical neurons with in the *LRRK2* R1441G mutation, baseline autophagy appeared to be enhanced.

**Figure 3.**
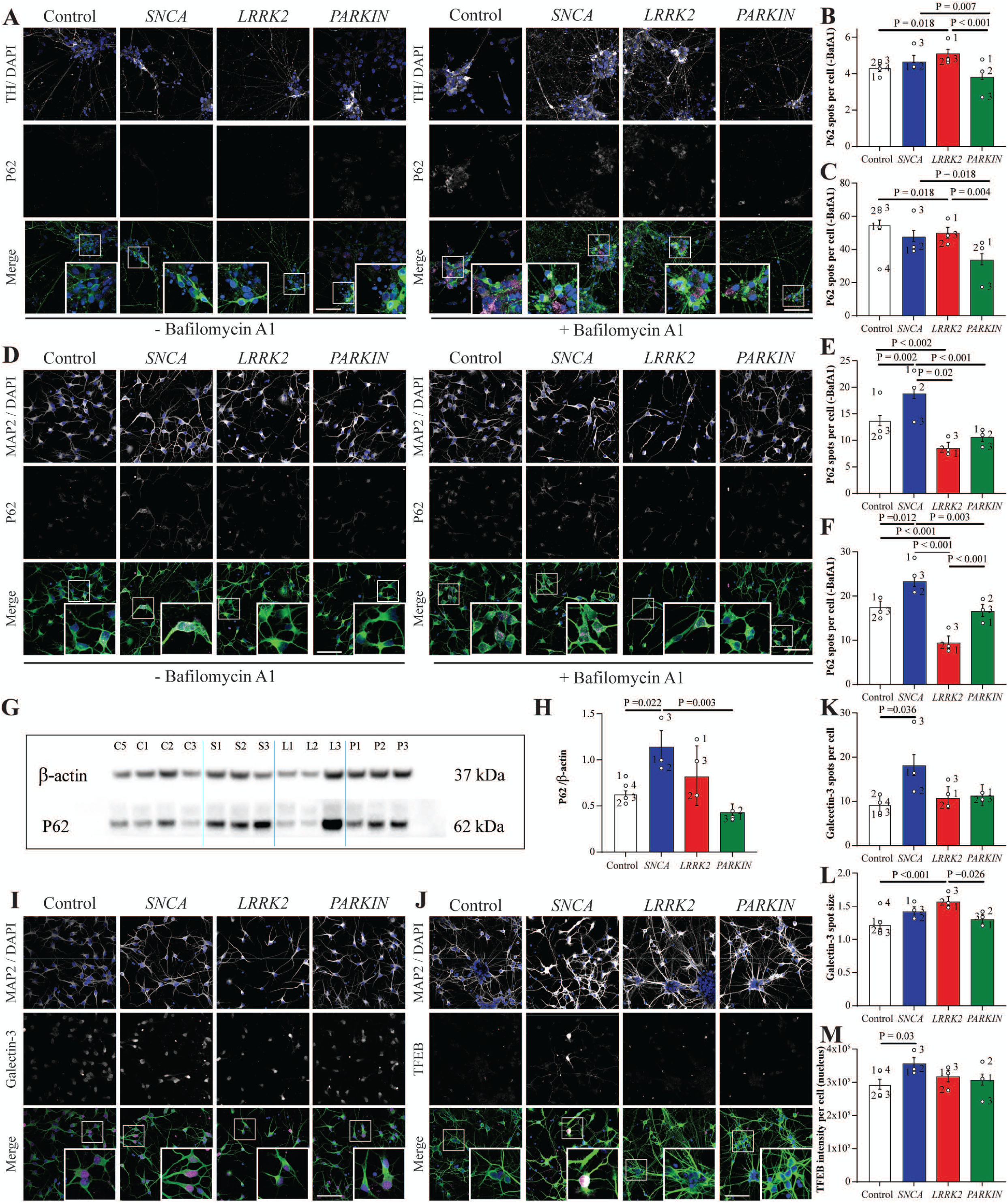
Autophagy markers in VMDA and cortical neurons show distinct phenotypes in *PRKN* and *SNCA* mutations. **A**) Representative images of VMDA neurons immunostained with P62 (Magenta), TH (green) and DAPI (blue) in the absence (left panel) and presence (right panel) of Bafilomycin A1. The number of P62 spots per cell in the different mutation groups of VMDA neurons in the absence (**B**) or presence (**C**) of Bafilomycin A1. **D**) Representative images of cortical neurons immunostained with P62 (Magenta), MAP2 (green) and DAPI (blue) in the absence (left panel) and presence (right panel) of Bafilomycin A1. The number of P62 spots per cell in the different mutation groups of cortical neurons in the absence (**E**) or presence (**F**) of Bafilomycin A1. Representative immunoblot (**G**) and subsequent quantification (**H**) of P62 protein levels in cortical neurons of different mutation groups. Protein expression is normalised to β-actin. **I**) Representative images of cortical neurons immunostained with Galectin-3 ((magenta), MAP2 (green) and DAPI (blue). **J**) Representative images of VMDA neurons immunostained with TFEB (magenta), TH (green) and DAPI (blue). The number (**K**) and size (**L**) of Galectin-3 spots in the different mutation groups of cortical neurons **M**) TFEB intensity per cell in the different mutauion groups of VMDA neurons. The scale bar in all images is 200 µm. Image quantification graphs show mean ± SEM. Each datapoint represents the mean value for each cell line derived from n=3 biological replications (each performed in at least duplicate).

### PRKN mutations are associated with impaired mitochondrial respiration in neuron types

Mitochondrial respiration was assessed in both neuron types as a measure of mitochondrial function. In the ventral midbrain cells, *PRKN* lof neurons exhibited a significantly reduced rate of oxygen consumption compared to control neurons (Fig. 4A), with both groups showing relatively low intra-cell line variability. While the oxygen consumption rate in *SNCA* A53T midbrain cultures was statistically lower than in controls, this effect was marginal (Fig. 4A), and the average values for *LRRK2* R1441G neurons were comparable to those of control neurons (Fig. 4A). However, both the *SNCA* and *LRRK2* mutation groups demonstrated high intra-cell line variability (Fig. 4A). In cortical neurons, the *PRKN* lof, *LRRK2* R1441G, and *SNCA* A53T mutation groups all exhibited significantly reduced rates of oxygen consumption compared to control neurons, with all groups showing relatively low intra-cell line variability (Fig. 4B). This consistency in oxygen consumption rates aligns with the stable percentage of mature cortical neurons observed in the cultures. A comparison between dopamine and cortical neurons revealed that *PRKN* neurons exhibited reduced oxygen consumption compared to controls in both cell types (Fig. 4A, B), with the effect being more pronounced in dopamine neurons, as quantified by regression analysis. Thus, *PRKN* lof impairs mitochondrial function regardless of neuron type, whereas *LRRK2* R1441G and *SNCA* A53T mutations have a more pronounced effect in cortical neurons.

**Figure 4.**
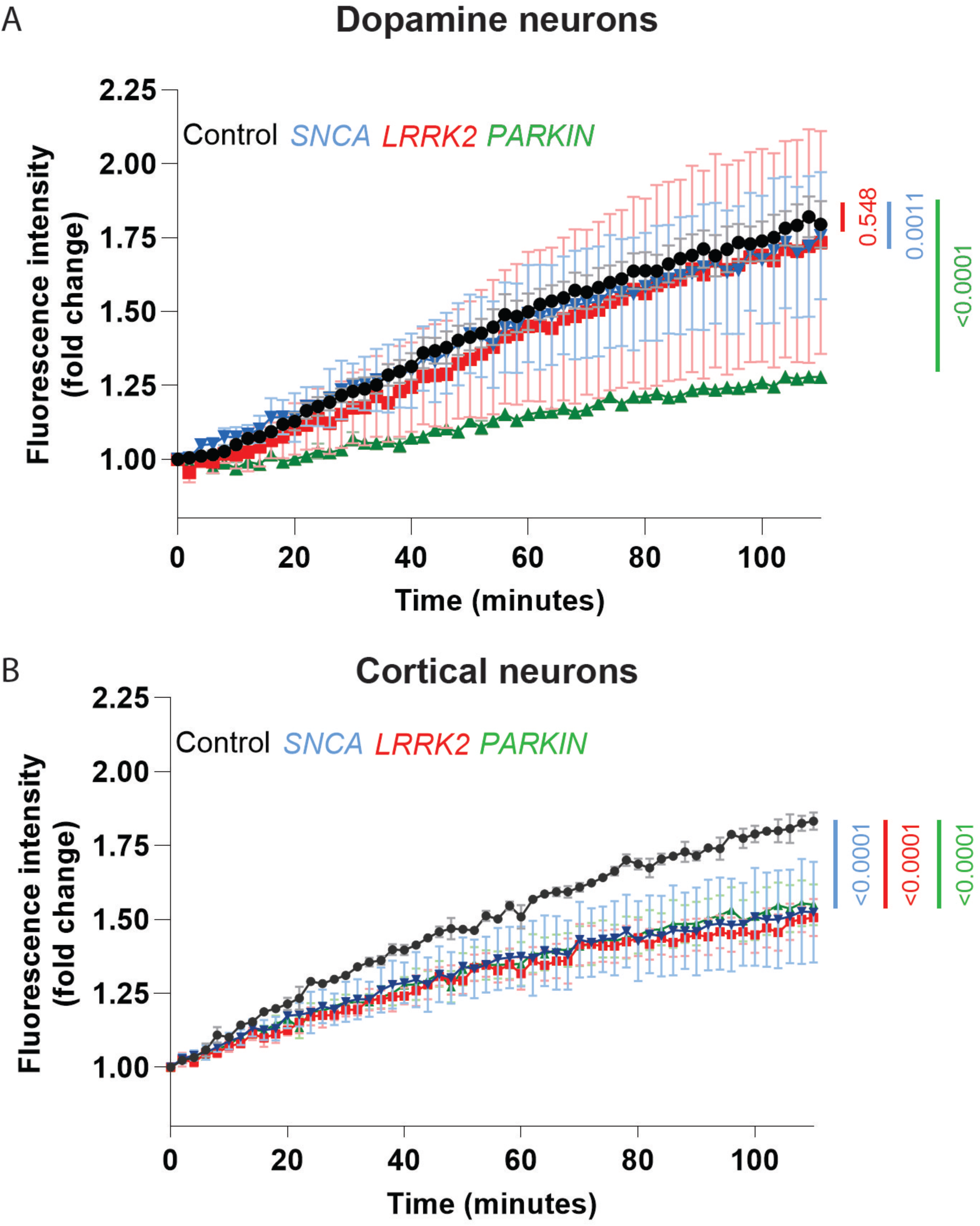
Mitochondrial respiration impairment in VMDA and cortical neurons with *PRKN, LRRK2* and *SNCA* mutations. Mitochondrial respiration was accessed by measuring extra cellular oxygen consumption rates of (A) dopamine neurons and (B) cortical neurons derived from control, *PRKN*, *LRRK2*, and *SNCA* mutation lines over a duration of 140 mins. Graphs show mean ± SEM at each timepoint. The slopes of PD groups were compared with the Control using One-Way ANOVA test, and their statistical significance levels were marked in the graph. Each datapoint represents the average of the three cell lines per mutation group, derived each from n=3 biological replicates.

### Alpha-synuclein pathology occurs in developing dopamine neurons with PRKN mutations

Alpha-synuclein pathology was assessed in the ventral midbrain dopamine and cortical neurons, noting that the staining intensity for the protein was much higher in dopamine cells than observed in cortical cells. For total alpha-synuclein, both the intensity and number of puncta per cell was increased in the TH+ *PRKN* lof cells (Fig. 5A-C). Whereas the intensity and number of Ser129 phosphorylated alpha-synuclein puncta were not changed across groups (Fig. 5D-F). In the cortical neurons, both the intensity of alpha-synuclein and the number of alpha-synuclein puncta per cell were significantly less in the *LRRK2* R1441G mutation (Fig. 5G-I). Despite weak signal, the number of phosphorylated alpha-synuclein puncta were significantly increased in the *LRRK2* R1441G mutation cells (Fig. 5J and L), whereas the intensity was not changed across groups (Fig. 5J and K). This data shows that PD mutations affect alpha-synuclein pathology in a cell and mutation type specific manner, with *PRKN* lof increasing alpha-synuclein puncta only in dopamine neurons, whereas *LRRK2* R1441G mutation had decreased numbers of alpha-synuclein puncta that were more phosphorylated at S129 only in cortical neurons.

**Figure 5.**
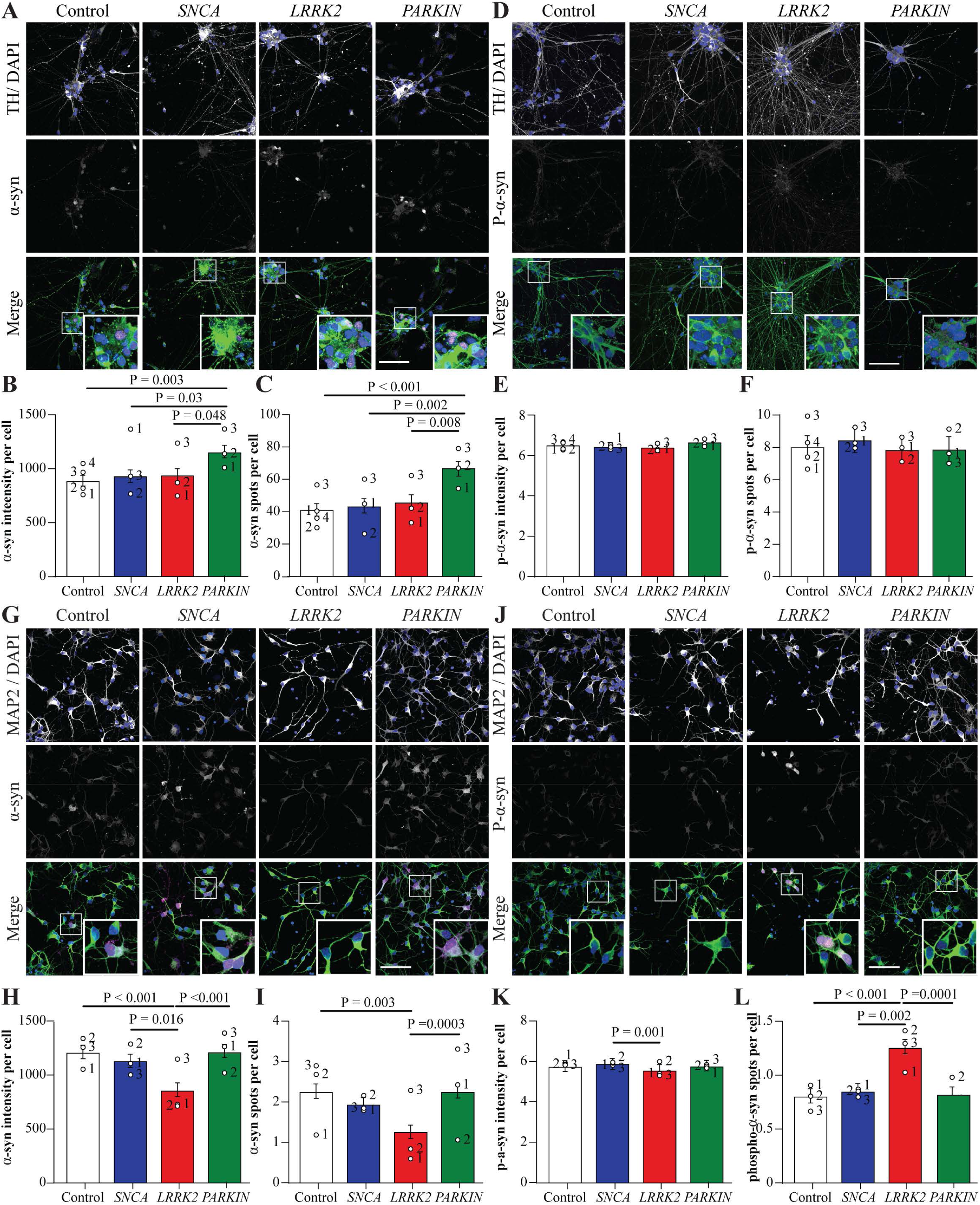
Alpha-synuclein staining shows distinct pathology in *PRKN* mutation dopamine neurons. **A**) Representative images of VMDA neurons immunostained with total α-sun (magenta), TH (green) and DAPI (blue). **B**) α-syn intensity per cell in the different mutation groups of VMDA neurons. **C**) Number of α-syn spots per cell in the different mutation groups of cortical neurons. **D**) Representative images of VMDA neurons immunostained with S129-phospho-α-syn (magenta), TH (green) and DAPI (blue). **E**) P-α-syn intensity per cell in the different mutation groups of VMDA neurons. **F**) number of p-α-syn spots per cell in the different mutation groups of VMDA neurons. **G**) Representative images of cortical neurons immunostained with α-syn (magenta), MAP2 (green) and DAPI. **H**) α-syn intensity per cell in the different mutation groups of cortical neurons. **I**) Number of α-syn spots per cell in the different mutation groups of cortical neurons. **J**) Representative images of cortical neurons immunostained with p-α-syn (magenta), MAP2 (green) and DAPI. **K**) P-α-syn intensity per cell in the different mutation groups of cortical neurons. **L**) Number of p-α-syn spots per cell in the different mutation groups of cortical neurons. The scale bar in all images is 200 µm. Image quantification graphs show mean ± SEM. Each datapoint represents the mean value for each cell line derived from n=3 biological replications (each performed in at least duplicate).

### Tau pathology occurs in developing dopamine neurons with LRRK2 and SNCA mutations

Levels of total and phosphorylated tau were assessed in both the dopamine and cortical differentiated neurons. For the dopamine neurons, the intensity of total tau was the same across groups, however the number of tau puncta per cell was significantly increased in both the *SNCA* A53T and *LRRK2* R1441G mutation groups (Fig. 6A-C). The intensity per cell of phosphorylated tau was increased in the *SNCA* A53T mutation cells, whereas the number of phosphorylated tau puncta were again significantly increased in both the *SNCA* A53T and *LRRK2* R1441G mutation groups compared to controls (Fig. 6D-F). In the cortical neurons, despite a trend for increased tau puncta in the *SNCA* A53T and *LRRK2* R1441G mutation cells (Fig. 6H and I), there was no significant effect of genotype for either the intensity per cell (Fig. 6G and H) or puncta per cell measures (Fig. 6G and H). There was also no change in the intensity per cell of phosphorylated tau (Fig. 6J and K), and unlike in dopamine neurons, tau puncta could not be robustly detected in the cortical neurons. This data shows that PD mutations affect tau pathology in a mutation type specific manner, with both *SNCA* A53T and *LRRK2* R1441G mutations having increased numbers of tau puncta in dopamine neurons, while *PRKN* lof did not display tau pathology.

**Figure 6.**
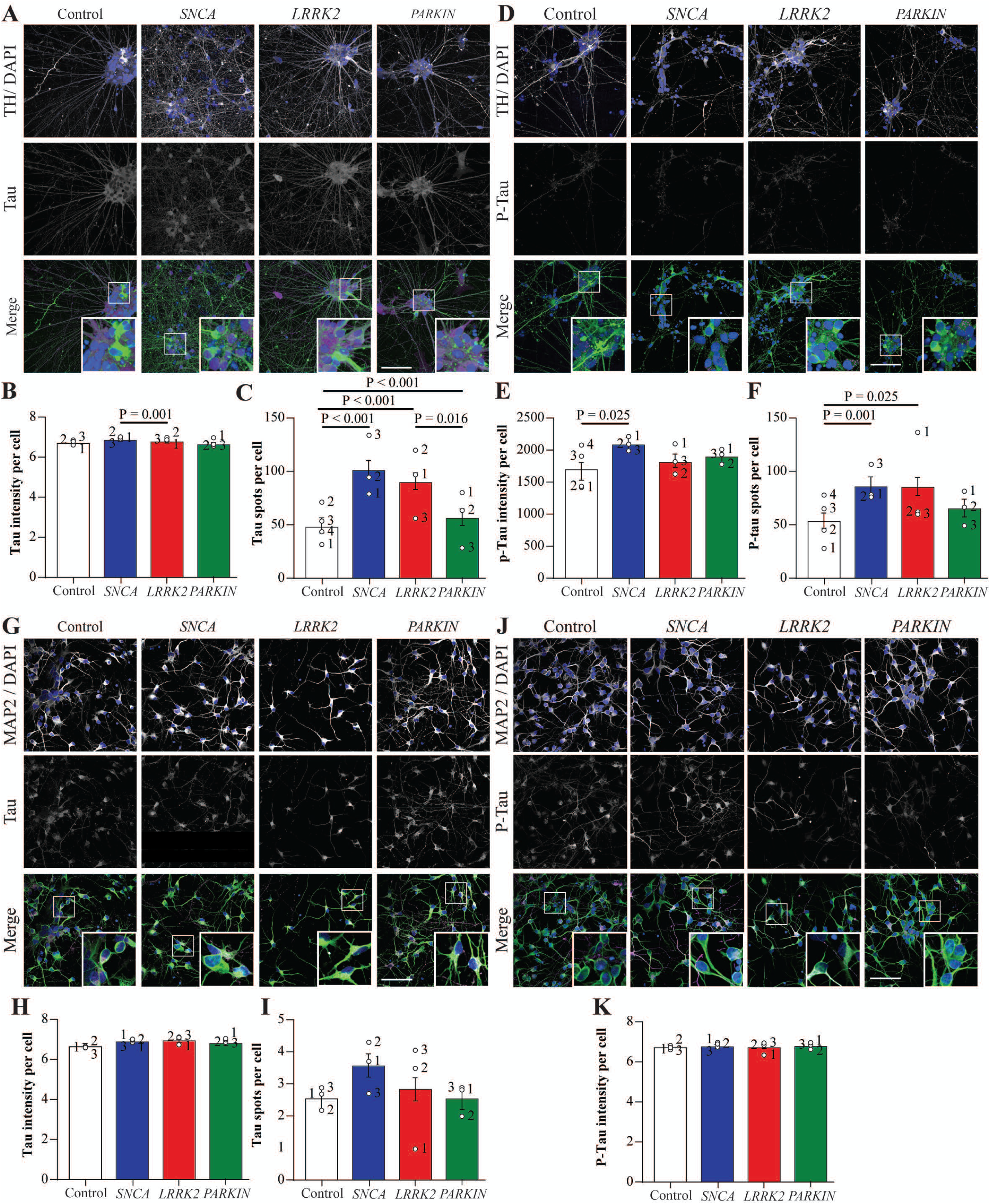
Tau staining shows distinct pathology in *LRRK2* and *SNCA* mutation dopamine neurons. **A**) Representative images of VMDA neurons immunostained with total Tau (magenta), TH (green) and DAPI (blue). **B**) Tau intensity per cell in the different mutation groups of VMDA neurons. **C**) Number of Tau spots per cell in the different mutation groups of cortical neurons. **D**) Representative images of VMDA neurons immunostained with p-Tau (magenta), TH (green) and DAPI (blue). **E**) P-Tau intensity per cell in the different mutation groups of VMDA neurons. **F**) number of p-Tau spots per cell in the different mutation groups of VMDA neurons. **G**) Representative images of cortical neurons immunostained with total Tau (magenta), MAP2 (green) and DAPI. **H**) Tau intensity per cell in the different mutation groups of cortical neurons. **I**) Number of Tau spots per cell in the different mutation groups of cortical neurons. **J**) Representative images of cortical neurons immunostained with P-Tau (magenta), MAP2 (green) and DAPI. **K**) P-Tau intensity per cell in the different mutation groups of cortical neurons. The scale bar in all images is 200 µm. Image quantification graphs show mean ± SEM. Each datapoint represents the mean value for each cell line derived from n=3 biological replications (each performed in at least duplicate).

## Discussion

The focus of this study was to compare PD mutations known to have divergent pathological phenotypes for evidence of cellular dysfunctions that may 1) underlie the very early selective vulnerability of dopamine neurons in *PRKN* lof mutations, and 2) underlie the pathological phenotypes. The cellular pathways assessed included lysosome and mitochondrial pathways in the selectively vulnerable dopamine neurons compared with cortical neurons which eventually accumulate pathological proteins without frank degeneration reported in post-mortem brain tissue. Three mutation types, *SNCA* A53T, *LRRK2* R1441G and *PRKN* lof, were selected as exemplars of divergent PD pathological phenotypes. Robust differentiation protocols were combined with a high content confocal imaging approach to compare phenotypes across the different cell lines.

### Cellular dysfunction selective to dopamine neurons occurred only in PRKN loss of function cell lines

The side-by-side comparison of different mutation types demonstrated that the most profound phenotypes in dopamine neurons were driven by *PRKN* lof. Dopamine neurons with *PRKN* lof showed a significantly greater reduction in mitochondrial respiration compared to control neurons, a result consistent across different studies employing *PRKN* lof lines [28-30]. *PRKN* lof dopamine, but not cortical neurons also had a significant reduction in lysosomal GCase activity and an inability to upregulate autophagy in response to bafilomycin. Lysosomal perturbations including impaired autophagy have been observed in *PRKN* KO iPSCs differentiated to dopamine neurons, including the same inability to upregulate p62 in response to bafilomycin [31]. Intriguingly, GCase activity was reported as increased in *PRKN* KO cells compared to isogenic control [31]. However, the *PRKN* KO cell study employed the 4-MUG GCase substrate that assesses glucosidase activity in whole cell lysates, as opposed to the PFB-FDglu probe used in the current study that specifically measures *in situ* lysosomal GCase activity. This could account for the differences observed as has been suggested [32]. Finally, cellular dysfunctions in *PRKN* lof dopamine neurons were found in occurrence with increased alpha-synuclein puncta. Although *PRKN*-associated PD is generally not associated with alpha-synuclein pathology, other studies of *PRKN* lof dopamine neurons have also found elevated levels of alpha-synuclein [33-35]. Although not consistently observed, increased alpha-synuclein has also been found in substantia nigra post-mortem brain tissue from PD patients with *PRKN* lof mutations [36], and at least one study has demonstrated that ∼60% of PD patients with a *PRKN* lof mutation may test positive on alpha-synuclein seeding amplification assay [37]. In contrast to dopamine neurons, studies of *PRKN* lof cortical cells are currently lacking. In this study, using *PRKN* lof cells from the same patients, it was intriguing that impaired mitochondrial respiration was also found, but there was no significant effect of *PRKN* mutation on measures of lysosomal function or alpha-synuclein puncta. These results appear consistent with the limited spread of alpha-synuclein pathology in brain tissue from *PRKN* lof mutation patients, and suggests that collective lysosomal and mitochondrial dysfunction specifically drives the selective degeneration observed in the more vulnerable dopamine neurons with *PRKN* lof mutations.

### Increased tau rather than alpha-synuclein occurred only in dopamine neurons in SNCA A53T and LRRK2 R1441G mutation cell lines

Examining the dopamine neurons, an interesting finding from this study is that both the *SNCA* A53T and *LRRK2* R1441G mutations were associated with a similar increased tau puncta phenotype, and increased phosphorylated tau puncta. Tau pathology has been linked to pathogenic *LRRK2* mutations before, including G2019S [38], I2020T [39] and R1441C [40] mutations. The *LRRK2* G2019S mutation has been reported to increase tau transmission [41] and LRRK2 kinase inhibitors have been reported to ameliorate pathology in *LRRK2* G2019S knockin mice seeded with tau fibrils [42]. Tau pathology has been less explored in the context of *SNCA* mutations, however increased tau pathology has been observed in postmortem brain tissue from *SNCA* A53T carriers [43, 44]. In the case of LRRK2, it has been suggested that defects in the autophagy pathway may affect tau clearance, and indeed the dopamine neurons with the *LRRK2* R1441G mutation had elevated levels of p62. While a trend was observed, levels of p62 were not significantly elevated in *SNCA* A53T dopamine neurons, as has previously been observed [45]. Moreover, levels of p62 were significantly elevated in *SNCA* A53T cortical neurons, but tau pathology was not evident. Thus, tau pathology seems unlikely to occur simply from impaired autophagy, and these mutations impact upon tau pathology via different mechanisms which require further investigation. In general, the observation that increased tau and not alpha-synuclein occurs early in dopamine neurons with these mutations supports the concept that tau may play a more central role in the selective degeneration of dopamine neurons in PD.

### Cellular dysfunction were dominant in cortical neurons and occurred in both SNCA and LRRK2 cell lines

Results from published studies are mixed on their read outs of alpha-synuclein levels in *SNCA* A53T lines, with some suggesting increased total alpha-synuclein levels, in particular compared to isogenic control lines [46-48], while others suggest an increase in early pathogenic species such as alpha-synuclein oligomers or phosphorylated alpha-synuclein [49, 50]. This is in contrast to numerous studies employing iPSC with alpha-synuclein multiplications, that consistently see a robust upregulation of alpha-synuclein phenotypes [45, 51-54]. Indeed, it has been suggested that the *SNCA* A53T mutation may drive pathology by altering alpha-synuclein aggregation propensity, rather than upregulation of protein levels themselves. In agreement, a recent study of iPSC differentiated to cortical neurons with the *SNCA* A53T mutation showed these cells have accelerated rates of alpha-synuclein seeding and oligomerization after inoculation with pre-formed fibrils [50]. This was coupled to increased levels of reactive oxygen species detected in the *SNCA* A53T cells [50]. In our study there was significantly reduced mitochondrial respiration in the cortical *SNCA* A53T cells, and further work could determine if this is linked to increased reactive oxygen species. Moreover, the *SNCA* A53T cortical neurons in our study also had increased levels of p62, suggesting impaired autophagy. Impaired autophagy can also contribute to accelerated seeding following inoculation of pre-formed fibrils [55] and this would be an area of further investigation. The data obtained on the *SNCA* A53T mutation lines supports the concept of cell type specific abnormalities that potentially impact on the development of cortical pathology over time.

The *LRRK2* R1441G mutation lines had a reduction in both mitochondrial respiration and lysosomal GCase activity in cortical neurons only. Other studies in cortical neurons with *LRRK2* mutations are lacking, but studies in dopamine neurons have shown reduced oxygen consumption with the *LRRK2* R1441C [19, 56], and G2019S [56] mutations. Why we did not also see mitochondrial dysfunction in dopamine neurons with the *LRRK2* R1441G mutation is unclear, however the variation between lines for this measure was very large and consequently our results are likely less conclusive. In contrast to dopamine neurons with *PRKN* lof mutations, the mitochondrial and lysosomal dysfunction in cortical neurons with *LRRK2* R1441G mutations did not associate with increased alpha-synuclein accumulation. Indeed, numbers of alpha-synuclein puncta were significantly decreased in the *LRRK2* R1441G lines, along with a reduction in p62. This may even suggest an improved clearance of alpha-synuclein in *LRRK2* R1441G cortical neurons. This is seemingly in contrast to several studies that have assessed *LRRK2* G2109S mutation cells differentiated to dopamine neurons, where there is a largely consistent upregulation of alpha-synuclein pathology [57-61]. However, it is increasingly being recognised that there are mechanistic differences between these two *LRRK2* mutation types. *LRRK2* G2019S increases kinase activity by ∼2-fold, whilst maintaining GTPase activity and 14-3-3 binding, whereas *LRRK2* R1441G increases kinase activity ∼5-fold, reduces GTPase activity and abolishes 14-3-3 binding [62-64]. Likewise, in studies of human brain tissue, *LRRK2* G2019S is associated with alpha-synuclein pathology in about 65% of cases, whereas the limited reports on R1441 mutations suggest this percentage is much lower [40]. In the current study there was also an increase in Ser129 phosphorylation of alpha-synuclein in the lower numbers of puncta in *LRRK2* R1441G cortical neurons, and thus further investigation as to why increased alpha-synuclein phosphorylation in combination with mitochondrial and lysosomal dysfunction paradoxically does not lead to alpha-synuclein accumulation, would be an important area of further investigation.

### Study limitations and future work

This study has several limitations to acknowledge. Although it is a large study in comparison to the current standard, sample sizes are still only three per group and expanding the sample size may allow for more subtle phenotypes to be uncovered. The inclusion of other mutations within the exiting genotypes (eg LRRK G2019S in addition to R1441G) or additional genotypes, for example *GBA1*, would also allow for further comparisons, and ultimately the inclusion of idiopathic lines may help to determine results in a broader context. Our assessment of tau and synuclein pathology was based on tools and reagents currently employed for postmortem diagnostic examinations, however, this could be extended to imaging for other pathogenic species using conformational specific antibodies or other phospho-epitopes. Also, we have only assessed cells at a single timepoint that is likely reflective of early events, and further neuronal maturation and/or time course studies could show different pathophysiological relationships at different disease stages. Finally, the developed high throughput analysis pipeline has enabled a comparative assessment of many neurons (of two differing regional specifications) from several iPSC lines and genotypes, and future studies employing additional techniques may contribute a more detailed understanding of the complex phenotypes uncovered.

## Conclusion

Overall, these findings demonstrate cell type specific dysfunctions with different PD mutations that are likely to impact on both selective neuronal vulnerability and the pathologies observed in PD. *PRKN* lof mutations selectively affected dopamine and not cortical neurons. While *PRKN* lof dopamine neurons had the expected significant reduction in respiration, they also had reduced lysosomal function (GCase activity) and impaired autophagy, all of which increased the levels of alpha-synuclein puncta. These selective changes were not observed in dopamine neurons from the other PD mutations or in *PRKN* lof cortical neurons, suggesting these early intrinsic cellular properties in dopamine neurons may drive early-onset *PRKN* disease. There were greater similarities between the *SNCA* and *LRRK2* mutations analysed, with both having increased tau puncta in their dopamine neurons, and mitochondrial and autophagy impairments in their cortical neurons (although the types of cortical cellular abnormalities were not the same). The cellular abnormalities in the *SNCA* A53T cell lines are likely to predispose to future alpha-synuclein aggregation, whereas in the *LRRK2* R1441G cell lines with increased alpha-synuclein phosphorylation, pathological accumulations were paradoxically not observed. Further analysis of these differences is warranted, particularly as cortical pathologies and symptoms are less likely with this *LRRK2* mutation in comparison to *SNCA* mutations (see www.mdsgene.org). That different PD-linked mutations drive cellular dysfunction to different extents across distinct neuron types will be important to consider for therapeutic development and stratification into clinical trials.

## Declarations

### Ethics approval

Experiments using human iPSCs were approved by the Human Research Ethics Committee at the University of Sydney (2017/094).

### Availability of data and materials

The data, code, protocols, and key lab materials used and generated in this study are listed in a Key Resource Table alongside their persistent identifiers at 10.5281/zenodo.13826736.

### Author’s contributions

Conceived the idea and secured funding – DK, GMH, CS, LT, CP, JJ

Planned the experiments – ND, GW, ALG, JC, CS, CP

Performed the experimental work – JC, GW, ALG, MZ, YL, DAB, CP, TF

Data analysis figures – ND, GW, JC

Drafted the manuscript – ND, GW, CS, GMH

Edited and approved the final version – all authors

## Acknowledgements

Data used in the preparation of this article were obtained from the Parkinson’s Progression Markers Initiative (PPMI) database (www.ppmi-info.org/access-data-specimens/download-data). For up-to-date information on the study, visit www.ppmi-info.org. The biospecimens and data for two PRKN cell lines used in this project were provided by NSLHD under approved protocol 0408-186M. We acknowledge the contribution of NSLHD to this research project. The authors acknowledge the Sydney Cytometry Core Research Facility, a joint initiative of Centenary Institute and the University of Sydney, for access to the Opera Phenix. We thank Heidi Cartwright for assistance with the preparation of figures.

