## Supplementary data for "Dopamine and cortical iPSC-derived neurons with different Parkinsonian mutations show variation in lysosomal and mitochondrial dysfunction: implications for protein deposition versus selective cell loss"

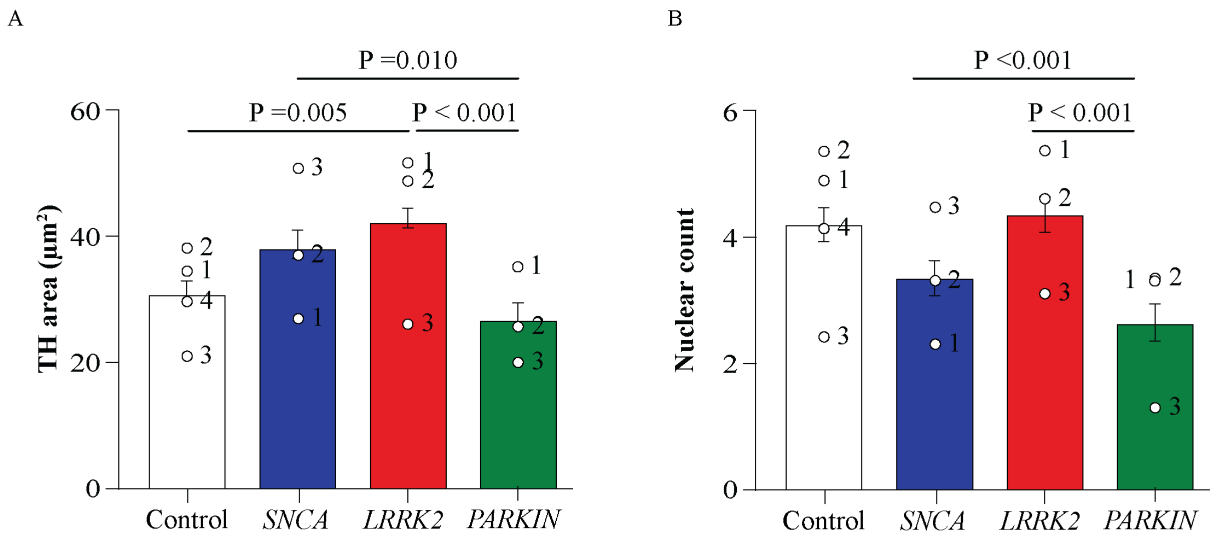


**Supplementary Figure 1. A**) Average TH area (µm2) calculated for each cell line of the different mutation groups in VMDA neurons. **B**) Nuclei count calculated for each cell line of the different mutation groups in VMDA neurons. Graphs show mean ± SEM. Each datapoint represents the mean value for each cell line derived from n=3 biological replications (each performed in at least duplicate).

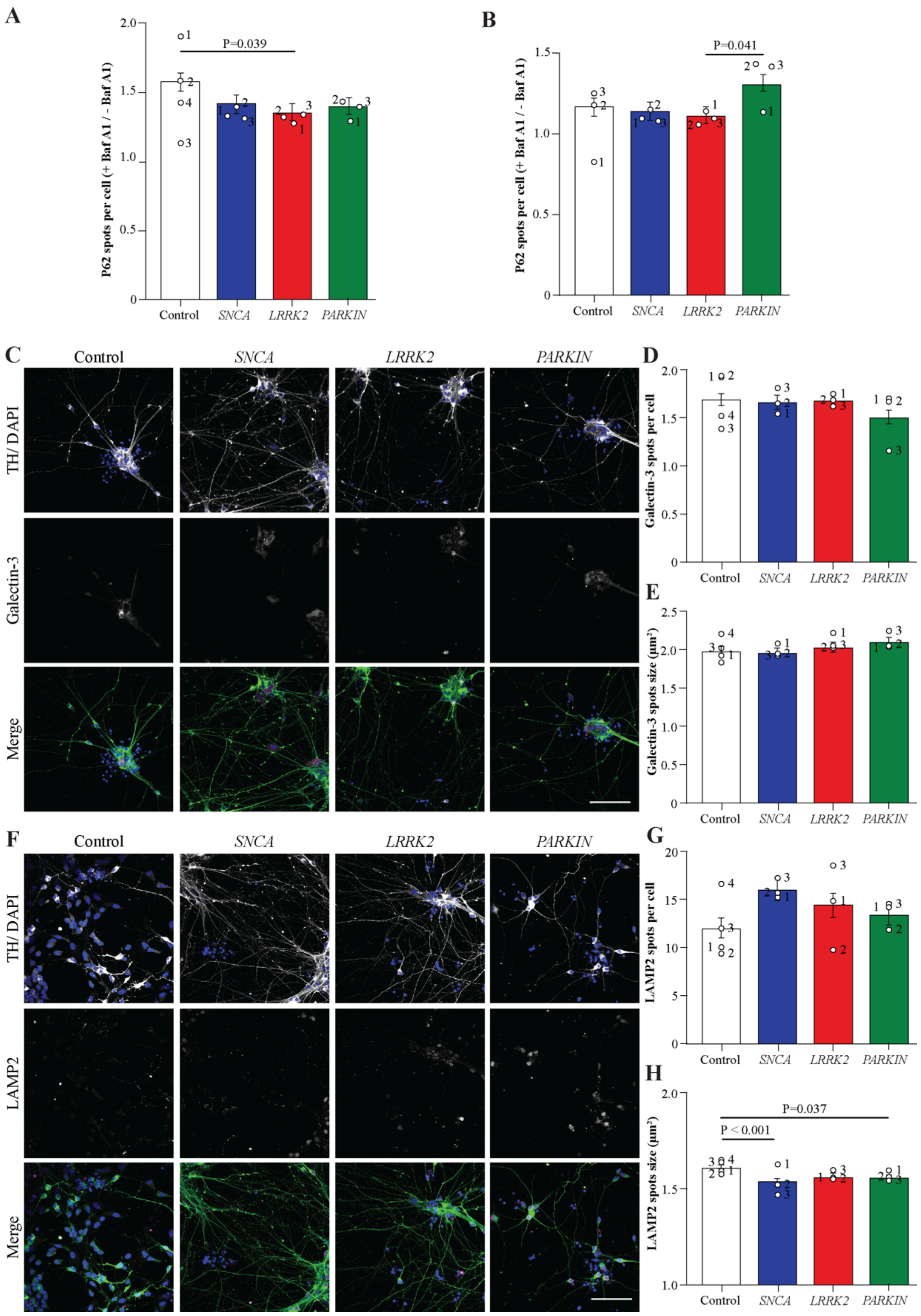


**Supplementary Figure 2.** **A**) The ratio of P62 spots in Bafilomycin A1 treated cells / non-treated for each cell line of the different mutation groups in VMDA neurons. **B**) The ratio of P62 spots in Bafilomycin A1 treated cells / non-treated for each cell line of the different mutation groups in cortical neurons. **C**) Representative images of VMDA neurons immunostained with Galectin-3 (magenta), MAP2 (green) and DAPI (blue). The number (**D**) and size (**E**) of Galectin-3 spots in the different mutation groups of VMDA neurons. **F**)

Representative images of VMDA neurons immunostained with LAMP2A (magenta), TH (green) and DAPI (blue). The number (**G**) and size (**H**) of LAMP2A spots in the different mutation groups of VMDA neurons The scale bar in all images is 200 µm. Graphs show mean ± SEM. Each datapoint represents the mean value for each cell line derived from n=3 biological replications (each performed in at least duplicate).

| Group | Cell line | RRID | Identifier | Age/sex | Disease | Mutation |
|---|---|---|---|---|---|---|
| **Control** | RM3.5 | NA | Control-1 | NA | - | - |
| | PPMI 11450-107 | CVCL_D5Q0 | Control-2 | 64/M | - | - |
| | PPMI 11302-101 | CVCL_D5NM | Control-5 | 76/M | - | - |
| | KOLF2-1J | CVCL_B5P3 | Control-3 | 55-59/M | - | - |
| **LRRK2** | PPMI 11576-101 | CVCL_D5SB | LRRK2-1 | 65/M | Unaffected | R1441G |
| | PPMI 14557-107 | CVCL_D5S9 | LRRK2-2 | 42/F | PD | R1441G |
| | PPMI 14555-107 | CVCL_D5RY | LRRK2-3 | 67/M | PD | R1441G |
| **PARKIN** | 01-060 C9 | CVCL_E3WU | PARKIN-1 | 58/F | PD | Ex3 del +int SNV |
| | 09-090 C18 | CVCL_D6I6 | PARKIN-2 | 39/F | PD | Ex2 del +ex2-5 del |
| | PK6 C2 / C4 | CVCL_C8H8 | PARKIN-3 | 49/M | PD | R275W + 132kb del |
| **SNCA** | PPMI 11555-104 | CVCL_D5SL | SNCA-1 | 36/M | PD | A53T |
| | PPMI 11557-103 | CVCL_D5S0 | SNCA-2 | 53/M | PD | A53T |
| | PPMI 11556-110 | CVCL_D5RU | SNCA-3 | 54/F | PD | A53T |

**Table S1. Cell line details.** Demographic and mutation information for cell lines used in this study. NA = not available.

### PRKN 09-090 C18

| Minimum iPSC Quality Control Panel | | | |
|---|---|---|---|
| **Genomic Baseline QC** | **Pass QC?** | **Method used** | **Results** |
| **Genotyping** | Yes | WGS | Exon deletion breakpoints confirmed |
| **Karyotype** | Yes | High-denity SNP array (Illumina GSA24) | Normal (part of the Lab resource manuscript submission to Stem Cell Research) |
| **Pluripotent phenotype** | Yes | PSC marker staining | iPSC colony morphology,nuclear OCT4/POU5F1, NANOG, manuscript submitted (Lab resource- Stem Cell Re |
| **Cell Line Identity** | Yes | STR (10 markers) | Cell line genetic identity with source fibroblasts confirmed |
| **Mycoplasma Testing** | Yes | MycoAlert ELISA | Data not shown |
| **Clonality** | Yes | double-picked, +10 passage screened | Data not shown |
| **Stability** | Yes | 30+ passage screened | Data not shown |
| **Extended iPSC Quality Control Panel** | | | |
| **Genomic Baseline QC** | **Pass QC?** | **Method used** | **Results** |
| **Copy Number Variation** | Yes | aCGH | All CNVs reported for each line |
| **Whole Genome Sequencing (can be performed with GP2)** | Yes | NGS | Available on request (Ryan Davis, USyd) |
| **p53 function or sequence** | Not analyzed | N/A | N/A |
| **BCL-2 Status** | Not analyzed | aCGH | N/A |

### PRKN 01-060 C9

| Minimum iPSC Quality Control Panel | | | |
|---|---|---|---|
| **Genomic Baseline QC** | **Pass QC?** | **Method used** | **Results** |
| **Genotyping** | Yes | WGS | Exon deletion breakpoints and point mutation confirmed |
| **Karyotype** | Yes | High-denity SNP array (Illumina GSA24) | No significant CNV or BAF deviations detected, resolution ~25kbps |
| **Pluripotent phenotype** | Yes | PSC marker staining | Phase-contrast iPSC colony morphology, positivity for nuclear OCT4/POU5F1, NNAOG |
| **Cell Line Identity** | Yes | STR (10 markers) | Cell line genetic identity with source fibroblasts confirmed |
| **Mycoplasma Testing** | Yes | MycoAlert ELISA | Data not shown |
| **Clonality** | Yes | double-picked, +10 passage screened | Data not shown |
| **Stability** | Yes | 30+ passage screened | Data not shown |
| **Extended iPSC Quality Control Panel** | | | |
| **Genomic Baseline QC** | **Pass QC?** | **Method used** | **Results** |
| **Copy Number Variation** | Yes | aCGH | All CNVs reported for each line |
| **Whole Genome Sequencing (can be performed with GP2)** | Yes | NGS | Available on request (Ryan Davis, USyd) |
| **p53 function or sequence** | Not analyzed | N/A | N/A |
| **BCL-2 Status** | Not analyzed | aCGH | N/A |

### PRKN 6 C4

| Minimum iPSC Quality Control Panel | | | |
|---|---|---|---|
| **Genomic Baseline QC** | **Pass QC?** | **Method used** | **Results** |
| **Genotyping** | Yes | WGS, Allele-specific Sanger sequencing | Missense variant and exon deletion confirmed |
| **Karyotype** | Yes | High-denity SNP array (Illumina GSA24) | Normal |
| **Pluripotent phenotype** | Yes | PSC marker staining, differentiation | Pavan…Ovchinnikov, Stem Cell Res DOI: 10.1016/j.scr.2023.103211 |
| **Cell Line Identity** | Yes | Allele Sanger haplotype sequencing | Cell line confirmed |
| **Mycoplasma Testing** | Yes | MycoAlert ELISA | Data not shown |
| **Clonality** | Yes | double-picked, +10 passage screened | Data not shown |
| **Stability** | Yes | 10+ passage screened | Data not shown |
| **Extended iPSC Quality Control Panel** | | | |
| **Genomic Baseline QC** | **Pass QC?** | **Method used** | **Results** |
| **Copy Number Variation** | Yes | aCGH | All CNVs reported for each line. |
| **Whole Genome Sequencing (can** | Yes | NGS | Available on request (Ryan Davis, USyd) |
| **p53 function or sequence** | Not analyzed | N/A | N/A |
| **BCL-2 Status** | Not analyzed | aCGH | N/A |

**Table S2. QC testing of new *PRKN* lof iPSC.** List of quality control tests performed on newly generated stem cell lines used in this publication. NA = not available

| Antibody | Company | Catalogue no. | Species | Dilution factor |
|---|---|---|---|---|
| BRN2 | Santa Cruz | Sc-6029 | Goat | 1:400 |
| TBR1 | Abcam | Ab31940 | Rabbit | 1:400 |
| CTIP2 | Abcam | Ab18465 | Rat | 1:400 |
| MAP2 | Thermofisher | PA1-10005 | Chicken | 1:500 |
| Total α-syn | BD biosciences | 610787 | Mouse | 1:400 |
| S129 phospho-α-syn | Abcam | Ab51253 | Rabbit | 1:300 |
| Total Tau (Tau5) | Novus Biologicals | NBP2-81091 | Rabbit | 1:300 |
| Phospho-Tau (AT8) | Thermofisher | MN1020 | Mouse | 1:300 |
| P62 / SQSTM1 | Abcam | Ab56416 | Mouse | 1:400 |
| Galectin-3 | Abcam | Ab2785 | Mouse | 1:400 |
| TFEB | Abcam | Ab267351 | Rabbit | 1:400 |
| LAMP2 | Abcam | Ab25631 | Mouse | 1:400 |
| FoxA2 | Thermofisher | H00003170-M01 | Mouse | 1:400 |
| TH | Thermofisher | PA1-5679 | Sheep | 1:500 |
| LC3B | Abcam | Ab192890 | Rabbit | 1:400 |

**Table S3. Antibodies.** Details for primary antibodies employed for immunocytochemistry and western blotting in this study.

| Antibody | Species | Company | Catalogue no. | Application | Dilution factor |
|---|---|---|---|---|---|
| Anti-Mouse IgG (H + L) HRP Conjugate | Goat | Bio-Rad | 1706516 | WB | 1:5000 |
| Anti-Rabbit IgG (H + L) HRP Conjugate | Goat | Bio-Rad | 1706515 | WB | 1:5000 |
| Anti-Mouse IgG Highly Cross-adsorbed Alexa Fluor 647 | Donkey | Thermofisher | A31571 | ICC | 1:500 |
| Anti-Mouse IgG Highly Cross-adsorbed Alexa Fluor 488 | Donkey | Thermofisher | A21202 | ICC | 1:500 |
| Anti-Mouse IgG Highly Cross-adsorbed Alexa Fluor 568 | Donkey | Thermofisher | A10037 | ICC | 1:500 |
| Anti-Rabbit IgG Highly Cross-adsorbed Alexa Fluor 488 | Donkey | Thermofisher | A21206 | ICC | 1:500 |
| Anti-Rabbit IgG Highly Cross-adsorbed Alexa Fluor 568 | Donkey | Thermofisher | A10042 | ICC | 1:500 |
| Anti-Chicken IgY (H + L) Highly Cross-adsorbed | Donkey | Sigma-Aldrich | SAB4600031 | ICC | 1:500 |
| Anti-Sheep IgG Cross Adsorbed Alexa Fluor 488 | Donkey | Thermofisher | A11015 | ICC | 1:500 |
| Anti-Rat IgG (H + L) Highly Adsorbed Alexa Fluor Plus 647 | Donkey | Thermofisher | A48272 | ICC | 1:500 |
| Anti-Goat IgG (H+L) Highly Cross-Adsorbed, Alexa Fluor Plus 647 | Donkey | Thermofisher | A32849 | ICC | 1:500 |

**Table S4. Secondary Antibodies.** Details for secondary antibodies employed for immunocytochemistry and immunoblot in this study.

| Stain / reagent | Company | Catalogue no. | Dilution |
|---|---|---|---|
| Cytopainter green (cytoplasmic stain) | Abcam | Ab176735 | 1:500 |
| DQ red BSA | Thermofisher | D12051 | 1:100 |
| PFB-FDGlu substrate | Thermofisher | P11947 | 1:1000 |

**Table S5. Live cell imaging probes.** Details for the live cell imaging probes employed in this study.

| | Measurement / Experiment | Magnification / Objective | Number of wells per plate per cell line | Z-stack: Number of planes | Z-stack: Step size | Number of fields of view imaged per well | Number of cells imaged per well average (SEM) | Channel/antibody combinations |
|---|---|---|---|---|---|---|---|---|
| **Live cell imaging** | **GCase activity: PFB-FDGlu expression** | 20x Water | 6 | 3 | 1.5 µm | 25 | 2382 (125) | Ch1: HOECHST Ch2: Alexa 488/PFB-FDGlu |
| | **Lysosomal activity: DQ-red BSA expression** | 40x Water | 3 | 3 | 0.5 µm | 25 | 720 (73) | Ch1: HOECHST Ch2: Alexa 488/CytoGreen Ch3: Alexa 555/DQ-red-BSA |
| **Fixed cells staining** | **Characterization: TBR1 expression** | 20x Water | 2 | 7 | 1.5 µm | 25 | 3898 (414) | Ch1: DAPI Ch2: Alexa 488/MAP2 Ch3: Alexa 555/TBR1 |
| | **Characterization: BRN2 and CTIP2 expression** | 20x Water | 2 | 7 | 1.5 µm | 25 | 3734 (373) | Ch1: DAPI Ch2: Alexa 488/CTIP2 Ch3: Alexa 555/BRN2 Ch4: Alexa 647/MAP2 |
| | **ALP marker: P62** | 40x Water | 6 | 10 | 0.5 µm | 49 | 1830 (145) | Ch1: DAPI Ch2: Alexa 488/MAP2 Ch3: Alexa 555/LC3B Ch4: Alexa 647/P62 |
| | **Pathology: α-synuclein + phospho-α-synuclein expression** | 40x Water | 3 | 8 | 0.5 µm | 49 | 2472 (197) | Ch1: DAPI Ch2: Alexa 488/MAP2 Ch3: Alexa 555/p-α-syn Ch4: Alexa 647/α-syn |
| | **Pathology: Tau 5 expression** | 40x Water | 3 | 8 | 0.5 µm | 49 | 1726 (147) | Ch1: DAPI Ch2: Alexa 488/MAP2 Ch4: Alexa 647/Tau5 |
| | **Pathology: phospho-Tau (AT8)** | 40x Water | 3 | 8 | 0.5 µm | 49 | 1748 (163) | Ch1: DAPI Ch2: Alexa 488/MAP2 Ch4: Alexa 647/AT8 |
| | **Lysosomal biogenesis and ruptured lysosomes: Galectin-3 and TFEB** | 40x Water | 3 | 10 | 0.5 µm | 49 | 1475 (199) | Ch1: DAPI Ch2: Alexa 488/MAP2 Ch3: Alexa 555/TFEB Ch4: Alexa 647/Gal3 |

**Table S6. Cortical neuron imaging parameters.** Details for the acquisition of images from cortical neurons.

| | Measurement / Experiment | Magnification / Objective | Number of wells per plate per cell line | Z-stack: Number of planes | Z-stack: Step size | Number of fields of view imaged per well | Number of cells imaged per well Average (SEM) | Channel/antibody combinations |
|---|---|---|---|---|---|---|---|---|
| **Live cell imaging** | **GCase activity: PFB-FDGlu expression** | 20x Water | 6 | 4 | 0.8 µm | 8 | 1236 (268) | Ch1: HOECHST<br>Ch2: Alexa 488/PFB-FDGlu |
| | **Lysosomal activity: DQ-red BSA expression** | 20x Water | 3 | 4 | 0.8 µm | 9 | 972 (151) | Ch1: HOECHST<br>Ch2: Alexa 488/CytoGreen Ch3: Alexa 555/DQ-red-BSA |
| **Fixed cells staining** | **Characterization: TH and FoxA2 expression** | 20x Water | 3 | 9 | 0.8 µm | 8-14 | 1404 (250) | Ch1: DAPI<br>Ch2: Alexa 488/MAP2<br>Ch3: Alexa 555/TH<br>Ch4: Alexa 647/ FoxA2 |
| | **ALP marker: P62** | 40x Water | 6 | 7 | 0.5 µm | 8-14 | 1623 (250) | Ch1: DAPI<br>Ch2: Alexa 488/TH<br>Ch3: Alexa 555/P62<br>Ch4: Alexa 647/LC3B |
| | **Pathology: α-synuclein + phospho-α-synuclein expression** | 40x Water | 3 | 10 | 0.5 µm | 8-14 | 1448 (189) | Ch1: DAPI<br>Ch2: Alexa 488/TH<br>Ch3: Alexa 555/α-syn<br>Ch4: Alexa 647/p-α-syn |
| | **Pathology: Tau 5 expression** | 40x Water | 3 | 10 | 0.5 µm | 8-14 | 2330 (430) | Ch1: DAPI<br>Ch2: Alexa 488/TH<br>Ch4: Alexa 555/Tau5 |
| | **Pathology: phospho-Tau (AT8)** | 40x Water | 3 | 10 | 0.5 µm | 8-14 | 928 (102) | Ch1: DAPI<br>Ch2: Alexa 488/TH<br>Ch4: Alexa 647/AT8 |
| | **Lysosomal biogenesis and CMA: Galectin-3 and LAMP2A** | 40x Water | 3 | 10 | 0.5 µm | 8-14 | 1312 (230) | Ch1: DAPI<br>Ch2: Alexa 488/TH<br>Ch3: Alexa 555/Gal3<br>Ch4: Alexa 647/LAMP2 |

**Table S7. Ventral midbrain neuron imaging parameters.** Details for the acquisition of images from ventral midbrain neuron cultures.
